# A Recurrent Neural Network for Rhythmic Timing

**DOI:** 10.1101/2024.05.24.595797

**Authors:** Klavdia Zemlianova, Amitabha Bose, John Rinzel

## Abstract

Despite music’s omnipresence, the specific neural mechanisms responsible to perceive and anticipate temporal patterns in music are unknown. To study potential mechanisms for keeping time in rhythmic contexts, we train a biologically constrained RNN on seven different stimulus tempos (2 – 8Hz) on a synchronization and continuation task, a standard experimental paradigm. Our trained RNN generates a network oscillator that uses an input current (context parameter) to control oscillation frequency and replicates key features of neural dynamics observed in neural recordings of monkeys performing the same task. We develop a reduced three-variable rate model of the RNN and analyze its dynamic properties. By treating our understanding of the mathematical structure for oscillations in the reduced model as predictive, we confirm that the dynamical mechanisms are found also in the RNN. Our neurally plausible reduced model reveals an E-I circuit with two distinct inhibitory sub-populations, of which one is tightly synchronized with the excitatory units.

## Introduction

The ability to estimate time is important for many abilities like dancing and playing or listening to a musical instrument. Experimental and computational studies of the neural mechanisms underlying the ability to estimate time have primarily focused on the temporal estimation of isolated intervals. The neural mechanisms underlying rhythmic timing remain unexplained.

Tapping along to an isochronous beat is the simplest form of beat-based timing and has been extensively studied in human psychoacoustic studies^1,2^. Recent work has shown that macaques, too, can be trained to synchronize their motor behavior to a beat^3,4^ although, unlike humans, they don’t spontaneously do so^5^. Electrophysiological recordings from macaque studies have characterized a number of features of neural data in rhythmic timing tasks that need to be explained by computational models including sequential firing patterns, ramping and cyclical patterns of neural activity, and stretching of neural firing patterns^3,6^. While previously proposed models for neural timing can show synchronization to a rhythmic stimulus and maintenance of the learned rhythm^7^, explaining the dynamical features of the observed neural data remains a challenge for existing models.

Motivated by recent successes of recurrent neural networks (RNNs) to reproduce dynamical features of neural data^8–12^, we train an RNN on a standard rhythmic timing task to replicate observed neural features from macaque experiments. Based on a neuro-meaningful context parametrization that mimics the driving current in a biophysical model, training leads to a controllable network that can produce oscillations over a wide range of frequencies and matches key dynamical features of experimental data. Using the RNN, we then develop a neurally plausible reduced model (E-I-I). Using this reduced model, we uncover dynamical mechanisms that explain the RNN’s oscillatory capabilities and propose distinct functional roles for two inhibitory subpopulations.

## Results

To examine potential mechanisms for rhythmic timing, we trained a recurrent neural network (RNN) with 500 units (80% excitatory, 20% inhibitory) on a synchronization and continuation task (SCT) (Fig. 1a) – a standard task in rhythmic timing literature. In this task, stimulus pulses are delivered equally spaced in time and a subject is asked to synchronize a motor action, such as a finger tap, with the stimulus sequence (synchronization phase) and then to continue producing that motor output at the same rate and phase after the stimulus is stopped (continuation phase). To model this task, the inputs to the RNN consisted of stimuli onset times (*I*_*Stim*_), modeled as short pulsatile inputs delivered at the frequency rate of the stimulus, as well as a context cue (*I*_*cc*_), modeled as a constant input drive whose amplitude was inversely related to the interval between stimulus pulses (Fig. 1b). The stimulus tones were presented for the first one second of network input only, while the context cue was provided for the entire two seconds of network input.

**Fig 1.**
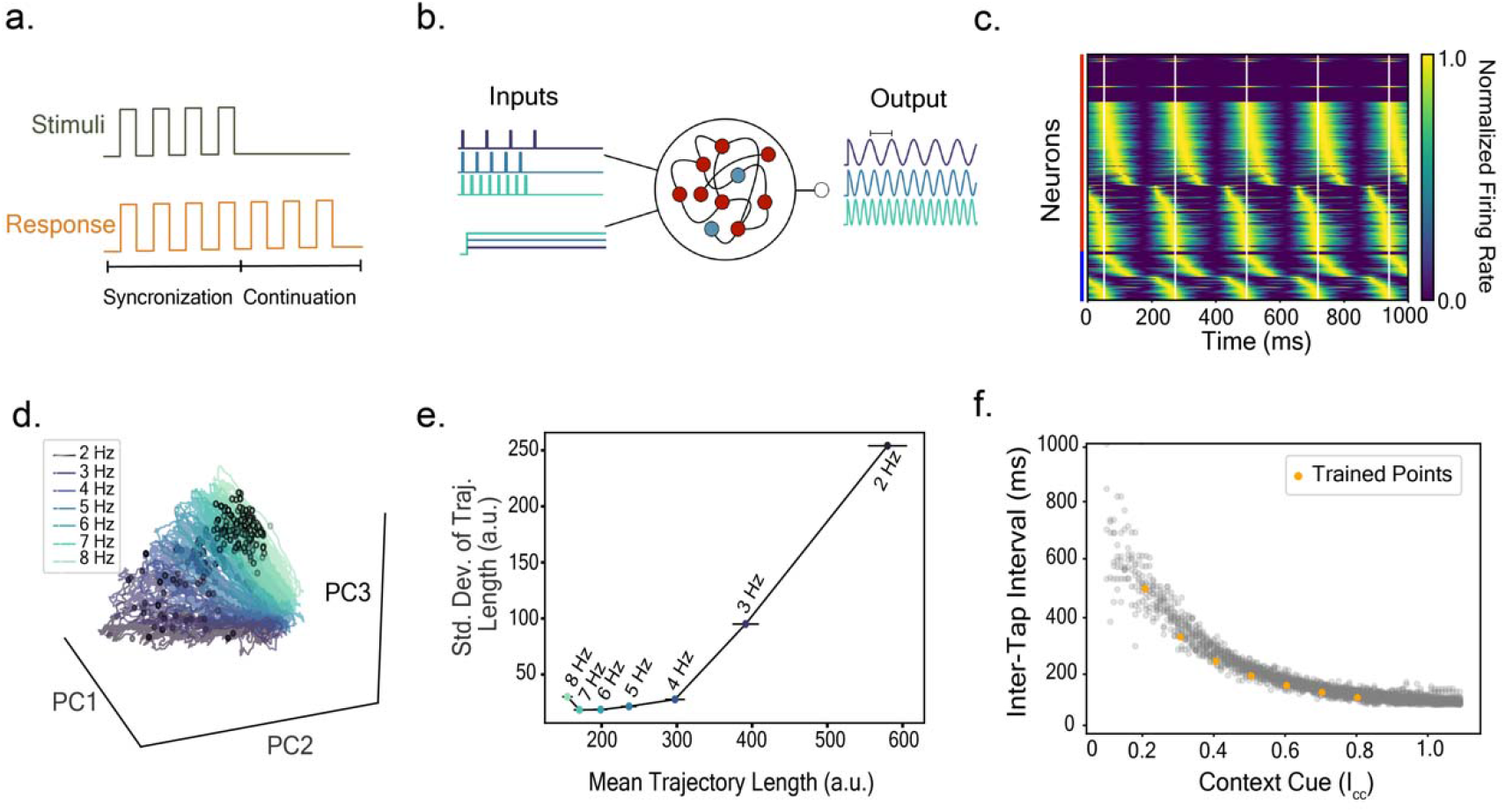
RNN trained on the Synchronization and Continuation Task replicates key features of neural dynamics. (**a**) Synchronization and continuation task. Stimulus pulses are presented equally spaced in time and the subject responds by first synchronizing their motor output with the stimulus (synchronization phase) and then continuing to produce the motor output at the same rate and phase in the absence of the stimulus (continuation phase). (**b**) Schematic of the RNN training. The RNN, composed of 500 hidden units with 80% excitatory (red) and 20% inhibitory (blue), receives two types of inputs: a context cue and stimulus pulses. The amplitude of the context cue is proportional to the frequency of the stimulus (purple = low frequency, teal =high frequency). The model output is a sinusoid of the same frequency as the stimulus input with the peaks of the sinusoids (“taps”) aligned to the stimulus tone times. Black horizontal bar on the output indicates one Inter-Tap Interval (ITI). (**c**) Firing rates of RNN units are normalized so that each unit’s activity is between 0 and 1 (color gradient) and then sorted by the time at which each unit hits its maximal firing rate during the first ITI. Excitatory (red) and Inhibitory (blue) units are sorted separately. White vertical lines indicate model “tap” times. (**d**) Firing time courses of RNN units trace out circles when projected to the space spanned by the top 3 PCs (explain 61% of the variance). Color gradient indicates frequency of stimulus (purple = low frequency to teal =high frequency). Black circles indicate model output “tap” times. (**e**) Mean length vs. standard deviation of trajectories from panel d. Distance is computed as the arclength of the discretized trajectory in ℝ^*N*^ taken between model output ‘tap’ times. Mean and standard deviation is computed across individual cycles for each *I*_*cc*_ (see Methods). (**f**) Simulated ITIs of the RNN output for *I*_*cc*_ (grey) at every 0.01 step from 0.1 to 1.1. Yellow circles indicate the points for which the RNN was trained. All dynamics for panels **c**.-**f**. are taken from the RNN simulated in the continuation phase, after transients have settled and the time course is nearly periodic, with noise *σ*_*in*_ = *σ*_*rec*_ = 0.01 (see Methods).

### RNN reproduces key features of experimental data

The SCT-trained RNN replicates several features of the neural dynamics previously observed in neural data of macaques performing the same task^3^. Sequential firing is a prominent feature of the neural recordings; different neurons fire during distinct phases on or between stimuli tones^3,6^. Furthermore, the same sequence of neurons fire in roughly the same order every timed interval. To see if our model replicates this observation, we perform similar analysis on the RNN hidden units (Fig. 1c). Since the peaks of the sinusoidal output of our model are a proxy for the tap events, we define the inter-tap-interval (ITI) for our model to be the time between consecutive peaks in the output. This analysis of the RNN hidden units reveals weak sequential activity that repeats each ITI (Fig. S1). This sorting also reveals that most excitatory units are synchronized and fire maximally around the “tap” times (white vertical lines in Fig. 1c) with some units having weaker firing peaks that lead or trail the tap-focused units. In contrast, inhibitory units show stronger sequential structure in their firing patterns with some units firing at maximum rates at different ITI phases. While the sequential activity is weaker than observed experimentally, the RNN replicates the over-representation of units firing at the “tap” just as observed in the experimental data^3^.

Examination of the experimental data also showed that when the neural firing rate time courses are projected onto the space spanned by the top three principal components (PCs), the trajectories trace out circles such that the radius and variability increase with stimulus period^3,6^. Projecting the firing rates of the RNN units into their corresponding space similarly reveal circular-like trajectories (Fig. 1d) that also increase in radius and variability (Fig. 1e) with stimulus period. The model ‘tap’ times (indicated by colored dots in Fig. 1d) align themselves along a line in state space - also in agreement with experimental findings.

When visualized in state space, the trained RNN trajectories are organized as a cone-like shape with slow trajectories (Fig. 1d dark colors, large radius) on one end and fast trajectories (Fig. 1d light colors, small radius) on the other end. Although we only trained the RNN on seven frequencies, the trained RNN could interpolate between the learned context cues (driving the RNN with intermediate values of *I*_*cc*_ produces oscillations of intermediate periods, see Fig. 1f) as well as extrapolate beyond the trained regime (Fig. 1f -- notice ITIs, shorter and longer, for contexts cues above 0.8 and below 0.2, respectively). The finding that the RNN is able to generalize is likely related to a low-dimensional structure in the network whose output varies continuously with a tonic context cue, consistent with previous studies^13–15^. However, unlike previous work on single-interval timing, our model’s generalization capabilities extend this finding to oscillatory systems. A context-parameterized structure is intuitively appealing as speeding up (slowing down) tempo can be accomplished by transitioning up (down) along the cone-manifold’s axis by increasing (decreasing) the neural drive (*I*_*cc*_).

### Developing a three-variable reduced model of the RNN

Having shown that the trained RNN reproduces several qualitative features of the neural data, we sought to understand the underlying dynamic mechanism of the RNN oscillatory solution by developing a reduced variable description. Close inspection of the RNN firing rate time courses (Fig. 1c) reveals that many of the units have highly similar activity patterns allowing us to identify four subpopulations: excitatory and inhibitory units that fire with a phase between (0.8,1) and (0, 0.2) of the “tap” time - which we will refer to as the tap excitatory (Tap-E) and tap inhibitory groups (Tap-I), respectively, as well as units that fire with a phase between 0.2 and 0.8 of the model “tap” times - the inter-tap excitatory (Int-E) and inter-tap inhibitory groups (Int-I), respectively (Fig. 2a). Simulations show that silencing the entire Int-E group in the trained RNN does not destroy the ability of the network to form oscillations, though it affects the frequency range of the network output (Fig. S2). Silencing any one of the remaining groups prevents the network from oscillating altogether suggesting that the remaining three groups are necessary to the mechanism.

**Fig 2.**
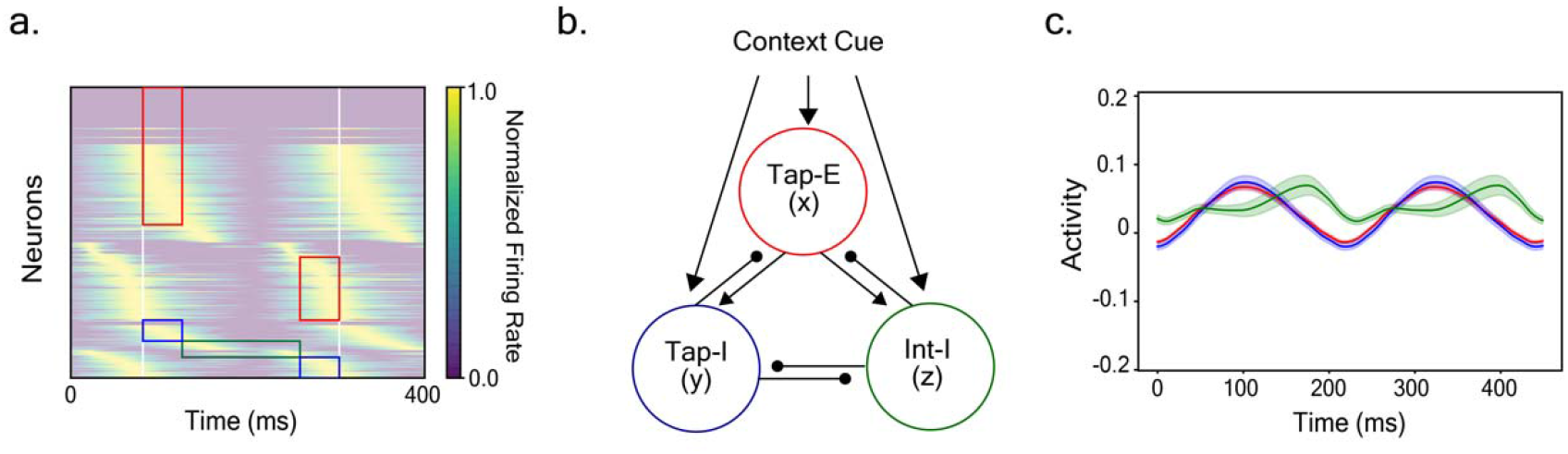
Distinct subpopulations in the RNN allow for the formulation of a three-variable reduced model. (**a**) Four distinct neuronal populations in the RNN are identified by first sorting the normalized firing activity (noise-free simulation) by peak firing time and grouping according to phase of peak firing rate. Example sorting during the continuation phase is shown for a stimulus of 5 Hz (*I*_*cc*_ = 0.5). Excitatory and inhibitory units are sorted separately. Excitatory and inhibitory units that peak within a phase of 0.2 to the tap times (white vertical lines) form the tap excitatory and tap inhibitory groups (red and blue boxes, respectively). Inhibitory units that have peak firing rates with phases between 0.2 and 0.8 form the inter-tap inhibitory (green box) group. (**b**) Schematic of reduced model with three populations of units defined in panel a. Connectivity weights between groups are taken to be group-averaged RNN connection weights except for a few weights (see Methods). (**c**) Time course of the pre-rectified activity (x-variables in equation 1 of Methods) of RNN hidden units (averaged over units) according to group membership: red, blue and green curves correspond to the average Tap-E, Tap-I andINT-I time courses, respectively. Time course shown during the continuation (no stimulus input) phase (*I*_*cc*_ = 0.5) and shading indicates ± standard error of the mean computed across units from the RNN simulated without noise during the continuation phase and after initial transients have settled, so that the individual units show their periodic response.

Motivated by this observation, we define a three-variable reduced model based on the three necessary RNN subpopulations : Tap-E (*x*), Tap-I (*y*), and Int-I (*z*) (Fig. 2b). The variables *x, y*, and *z* correspond to the subgroup-averaged activities of the hidden units.

The evolution of these variables is defined by rate equations modeled after the formulation of the RNN:

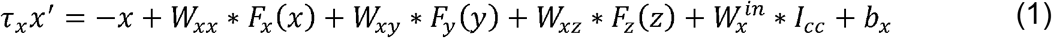

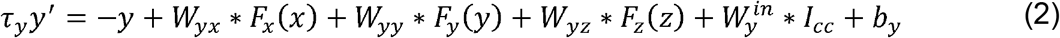

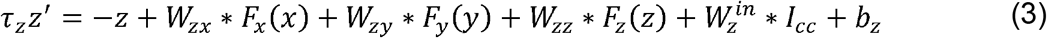

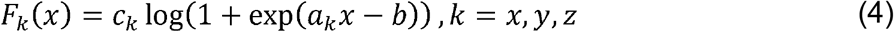

where connectivity weights between units and the input weight on the context cue (*I*_*cc*_) are given by *W* and *W*^*in*^, respectively. The bias and time constant for each population *k* are given by *b*_*k*_ and *τ*_*k*_, respectively Since we are interested in the structure of the model that maintains oscillations during the continuation phase, we drop the stimulus tone input so that the only external input to the reduced model is the context cue (*I*_*cc*_). To approximate the effects of the RNN units’ heterogeneous properties, we replace the non-linearity in the reduced form with a soft-plus function, *F*(·), that is fit to the averaged ReLU functions for each subpopulation (equation (4)). Finally, we take the connectivity weights to be the group average weights from the RNN except for *τ*_*z*_, *W*_*zy*_, *W*_*zz*_ which we have hand-tuned (see Methods). The reduced model is simulated with zero noise, throughout.

**Table 1:** Reduced Model Parameters.

| Parameter | Value |  |  |
| --- | --- | --- | --- |
| $W_{xx}$ | 8.949 | $b_z$ | -0.1 |
| $W_{xy}$ | -7.04 | $\tau_x, \tau_y$ | 10ms |
| $W_{xz}$ | -2.952 | $\tau_z$ | 50ms |
| $W_{yx}$ | 9.123 | $a_x$ | 150.134 |
| $W_{yy}$ | -7.386 | $b_x$ | -0.476 |
| $W_{yz}$ | -3.072 | $c_x$ | 0.007 |
| $W_{zx}$ | 8.935 | $a_y$ | 62.873 |
| $W_{zy}$ | -4 | $b_y$ | 0.481 |
| $W_{zz}$ | -1 | $c_y$ | 0.016 |
| $W_x^{in}$ | 0.048 | $a_z$ | 87.0 |
| $W_y^{in}$ | 0.055 | $b_z$ | -0.781 |
| $W_z^{in}$ | 0.068 | $c_z$ | 0.012 |
| $b_x, b_y$ | 0 | | |

As a first step, we look at the subgroup-specific activity of the RNN hidden units. Averaging the hidden unit activity in the RNN according to subgroup (Fig. 2c) revealed low amplitude activity for all three groups. Furthermore, the Tap-E and Tap-I groups appeared to be highly synchronized and nearly identical while the Int-I group showed a phase offset in its time of peak firing compared to the Tap groups.

### Analysis of the three-variable model

Qualitatively, we see agreement between the dynamics of the reduced model and the RNN. For small and large values of context cue (*I*_*cc*_), the system converges to a steady state (Fig. 3a and Fig. 3c, respectively). For intermediate values of *I*_*cc*_, the reduced model displays oscillations (Fig. 3b) with *x* (red) and *y* (blue) variables highly synchronized and *z* (green) showing a phase delay, in agreement with the RNN behavior of the averaged Tap-E, Tap-I and Int-I time courses shown in Fig. 2c. The RNN dynamics also converge to a steady state for *I*_*cc*_ values beyond the RNN’s extrapolation regime, *I*_*cc*_ < 0 and *I*_*cc*_ > 1.1 (not shown). The reduced model produces a wide range of oscillation frequencies spanning 1 to 17Hz (Fig 3d) which overlaps with the produced frequency range of the RNN (Fig. 1f). The rise of ITI with decreasing *I*_*cc*_ is steep and localized rather than gradual as in the RNN; the difference is perhaps lack of mechanisms for sequential activity or explicit account of heterogeneity in the reduced model.

**Fig 3.**
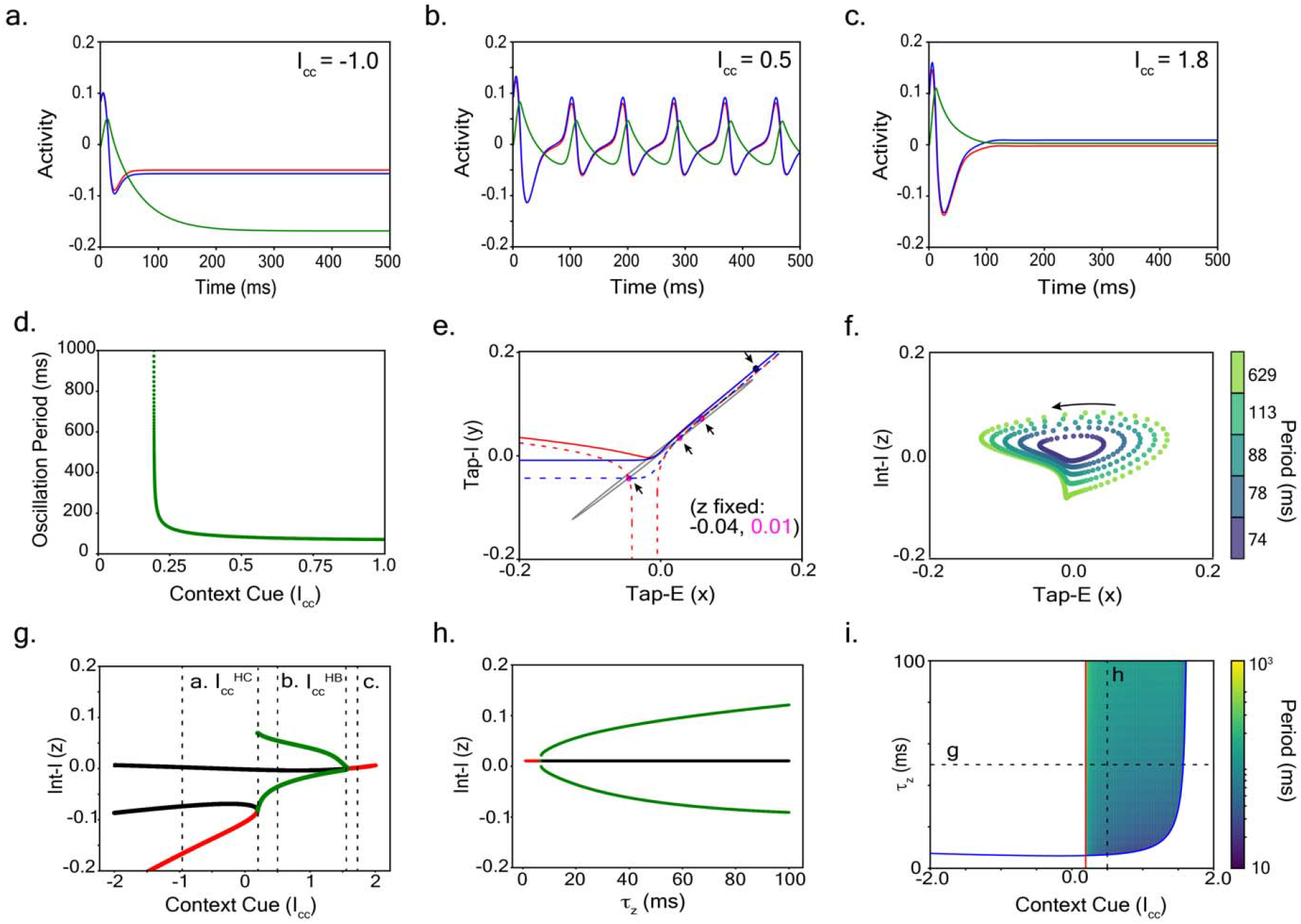
In the reduced model, long-period oscillations arise from a homoclinic bifurcation and disappear as short-period oscillations via a Hopf bifurcation. (**a**) Example time courses of *x* (red), *y* (blue), and *z* (green) variables reduced model (equations (1)-(4)) simulated with context cue *I*_*cc*_ = −1.0 and *τ*_*z*_ = 50*ms*. (**b, c**) Time courses of the reduced model for context cues *I*_*cc*_ = 0.5 and *I*_*cc*_ = 1.8. Colors are the same as in **a**. (**d**) Oscillation period vs. context cue for the reduced model shown for *τ*_*z*_ = 50*ms*. (**e**) Two-dimensional phase plane projection of reduced model for *x* vs. *y*.Nullclines for *x* (red) and *y* (blue) variables with *z* fixed at *z =* − 0.04 and *z =* 0.01 (solid and dashed lines). Arrows indicate locations of nullcline intersection points for *z =* − 0.04 (one point) and *z =* 0.01 (three points) which are also shown in black and magenta circles, respectively. Example model trajectory for *I*_*cc*_ = 0.5 shown in grey. (**f**) Example oscillatory trajectories for context cue values *I*_*cc*_ = 0.197, 0.3, 0.5, 0.8 and 1.2 (gradients from yellow to purple; dots plotted for every 1 *ms* timestep) projected into the two-two-dimensional phase plane, *z* vs. *x* Direction of flow is counterclockwise (black arrow). (**g**) One dimensional bifurcation diagram of the reduced model showing the effect of context cue (*I*_*cc*_) on model dynamics. The homoclinic and Hopf bifurcation points are indicated by 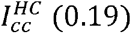 and 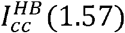, respectively. Red and black lines indicate stable and unstable fixed points, respectively. Green lines indicate the max and min z-values of the stable periodic orbit. Dashed lines at context cue levels of *I*_*cc*_ = −1, *I*_*cc*_ = 0.5 and *I*_*cc*_ = 1.8 correspond to panels a, b, and c, respectively. (**h**) One-dimensional bifurcation diagram with *τ*_*z*_ as the control parameter. Hopf bifurcation occurs at *τ*_*z*_ = 6.6*ms*. Colors same as in g. (**i**) Two-parameter bifurcation diagram for control parameters context cue (*I*_*cc*_) and time constant *τ*_*z*_. Homoclinic bifurcation point 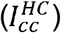 and Hopf bifurcation point 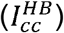 as a function of context cue and *τ*_*z*_ indicated by red and blue lines, respectively. Color indicates period of oscillations in log-scale (see color bar). White areas are those in which no stable oscillations exist. Dashed lines indicate the slices shown in panels g and h.

To understand the properties of the reduced model, we look at various two-dimensional projections of the phase space at *I*_*cc*_ = 0.5, for which there is an oscillatory solution in the reduced model (Fig. 3b). We consider snapshots of the nullclines at particular times with the third variable fixed (full motion of the nullclines and dynamics can be seen in Movie 1). Motivated by the fact that *z* evolves more slowly than *x* and *y* (since *τ*_*z*_ is large relative to *τ*_*x*_ and *τ*_*y*_), we chose two different projections for small and large values of *z*.In the *x* vs. *y* plane (Fig 3e), we plot the projection of two different *x*-nullclines (red) and the *y*-nullcline (blue) for a fixed large (dashed) and small (solid) value of *z*. For small *z* (0.04), note that the solid nullclines largely overlap and are nearly linear for *x, y* > 0 (we informally refer to the region of balance where *x* ≈ *y* as the balance manifold) with exactly one intersection point (black circle in Fig. 3e). The projection of the oscillatory solution onto this phase plane when is small lies near this linear portion, suggesting that the *x* and *y* variables (representing Tap-E and Tap-I inhibitory populations) are balanced closely with one another. This portion of the trajectory corresponds to the red and blue boxed regions of Fig. 2a. For larger *z* (0.01, dashed), the nullclines retain the overlap in the linear region, but show a large change in shape of the nullcline for *y* < 0, which now intersect at three points indicated by magenta circles in Fig. 3e. These intersection points for both values of *z* are not actual fixed points of the three-three-dimensional system since *z* varies over the course of the oscillation. However, they do transiently affect the dynamics. In particular, the bottom intersection point (where *x, y* < 0) transiently acts like a stable attracting point, causing the projection of the oscillatory trajectory to pass through a neighborhood of this point. Since *z* evolves relatively slowly, the *x* and *y* nullclines change slowly thus keeping the trajectory near this bottom intersection point for some time. The result of this can be seen in Fig. 3b where the *x* and *y* time courses have a plateau region around 0. As *z* is decreased, the 0 disappears allowing the trajectory to move towards the *x, y* > 0 region. The projection of the periodic solution onto the *x* vs. *z* plane (Fig. 3f) shows how the amplitude and period of the solution changes as a function of the context cue (*I*_*cc*_). Observe that points along each trajectory are plotted at each 1*ms* increment, implying that longer period trajectories slow down near the south end of the trajectory.

Given that both the *z* variable and *I*_*cc*_ enter additively in equation (1), we asked whether changes in *I*_*cc*_ would impact the position of *x* and *y* nullclines in a manner similar to how *z* affects the nullclines. We performed a bifurcation analysis using *I*_*cc*_ as the control parameter and report the results in Fig. 3g. For *I*_*cc*_ negative there can be multiple steady states but in this parameter regime, only one is stable (stable in red, unstable in black). Since *x, y*, and *z* are negative for this stable state, this corresponds to a state of no firing. There is a single stable state for large enough *I*_*cc*_ (Fig. 3g red line for *I*_*cc*_ > 1.57). Oscillations exist for intermediate *I*_*cc*_ trough to peak amplitude (Fig 3g, green for max and min of oscillation during a cycle) decreasing and frequency increasing as *I*_*cc*_ increases.

To understand how oscillations arise and disappear in this system, we take a closer look at the how the fixed points change stability. Starting at small values of the context cue (*I*_*cc*_ < 0.19), there are three fixed points whose existence can be seen in Fig.3g (black and red lines). As the context cue increases, the lower two branches of the bifurcation curve meet at a saddle node point and disappear. This is the parameter value at which the system changes between a regime with a stable steady state attractor to an oscillatory regime. The oscillation that appears for 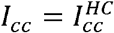 has large amplitude and zero frequency (infinite period); it’s called a homoclinic orbit, a unique long for *I*_*cc*_ near to but just greater than 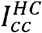 and decreases with increasing *I*_*cc*_ . At large values of context cue (*I*_*cc*_ > 1.57), there is only one fixed point and it is a stable attractor. As the context cue is decreased from the right side of the Fig. 3g, oscillations emerge with small amplitude by destabilizing the steady state via a Hopf bifurcation 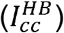. These dynamical features reveal that for context cues less than 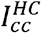, the reduced model converges to a stable fixed point (e.g., Fig 3a); for context cues between 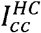 and 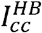, the system shows stable oscillations (e.g., Fig 3b); and for context cue values above 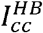, the system again converges to a stable fixed point via a spiraling attractor (e.g., Fig 3c). Furthermore, given that the reduced model is meant to inform the RNN, these observations allow us to make the prediction that for the RNN, as the context cue increases, oscillations arise as long period, large amplitude oscillations from a homoclinic bifurcation and disappear as short period, small amplitude oscillations via a Hopf bifurcation.

In the RNN, we already observed that Tap-E and Tap-I are in balance while the Int-I is phase delayed (Fig. 2c) thereby acting like delayed inhibition. Slow feedback inhibition is a known way to create oscillations in many systems and motivated by this, we explore how the *z* variable affects oscillations in our reduced model. We quantify this with a bifurcation analysis using *τ*_*z*_ as a control parameter. With fixed context cue *I*_*cc*_, the reduced model transitions from a stable steady state to stable oscillations via a Hopf bifurcation (Fig. 3h). If the *z* variable is too fast, the system does not oscillate. This result for the reduced model suggests that in order for the RNN to oscillate, there must exist a sufficiently long window of inhibition. Recalling that the time constants of the Int-I units are short (10 *ms*), the sequential firing patterns of Int-I is what establishes the sufficiently long window of inhibition for the RNN to oscillate. However, at *τ*_*z*_ = 10*ms*, the reduced model’s oscillation period for a given *I*_*cc*_ is much shorter than that of the RNN. Therefore, to obtain a comparable range of frequencies to the RNN, we chose a longer *τ*_*z*_ (50 *ms*).

### Assuming E-I balance allows further reduction to a two-variable model

The tight balance observed between the Tap-E (*x*) and Tap-I (*y*) variables suggests a further reduction of our model to a two-variable system by assuming that *x* and *y* are related by a scalar multiple taken to be the slope of the overlapping region of the nullclines (Fig. 3a solid line *x, y* ≥0). Introducing this assumption (*y=*1.1*x*), the resulting two-variable system (Fig. 4a) is amenable to phase plane analysis without the need for projecting from a higher dimensional space. The oscillation trajectory strikingly resembles the projected limit cycle of the *x* − *y* − *z* model (compare Fig. 4a to 3f). An example trajectory is plotted at every 1 *ms* timestep (grey) and shows a slowing down near the southern end of the trajectory as observed in the three-variable reduced model. Furthermore, the qualitative dynamical features are preserved: oscillations emerge with long period via a homoclinic bifurcation that then disappear via a Hopf bifurcation as *I*_*cc*_ changes from medium to high values (compare Fig. 4b to 3g). Finally, we see that the system retains small amplitude oscillations and a phase delay in the peak activity of the *z* variable (compare Fig. 4c to 3b).

**Fig 4.**
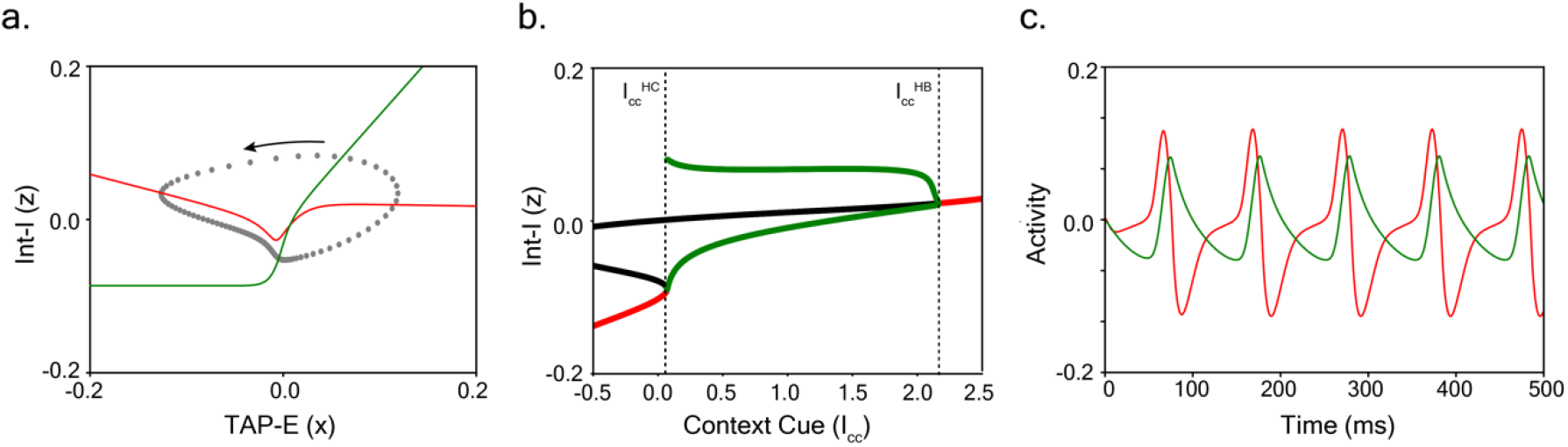
Two variable reduced model. **a**. Phase plane representation of the two-variable reduced model (obtained by assuming strong E-I balance: *y* = 1.1 *x*) for Tap-E (*x*) and Int-I (*z*) variables. The limit cycle trajectory (grey; dots plotted for every 1 *ms* of trajectory) is a global attractor, *I*_*cc*_ = 0.2; it crosses the *x*-nullcline (red) vertically and the *y*-nullcline horizontally (green). The flow is counterclockwise (black arrow); peak of the Int-I inhibition occurs on the downstroke of the Tap-E excitation. **b**. Bifurcation diagram of the two-variable reduction with context cue (*I*_*cc*_) as control parameter resembles that for three-variable model (Fig. 3e). The limit cycle attractor (green) transitions to a stable steady state (red) via homoclinic and Hopf bifurcations for 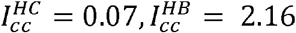, respectively. Black lines indicate unstable fixed points. **c**. Example time course of *x* and *y* variables for the same trajectory shown in **a**.

### Analysis of RNN dynamics confirms predictions from reduced model

Having identified the reduced models’ dynamic structure and attractor transitions, we searched for the corresponding features in the RNN. We identified a two-dimensional plane that captures the key fixed points affecting the RNN dynamics and investigated the changes in the RNN dynamics as a function of changes in context cue, (*I*_*cc*_) in this plane. To identify the relevant plane, we performed Principal Component Analysis on RNN time courses from trajectories initialized on the balance manifold in the non-oscillatory regime (see Methods) and chose the plane spanned by the first two PCs (this captures 80% of the variance for all noise-free RNN trajectories). Note that the projection plane was kept constant for the following analysis.

First, consider the RNN behavior for context cue values for which the system converges to a unique stable steady state. The top row of Fig. 5a shows the projection of three example trajectories (red curves) for three small values of the context cues (*I*_*cc*_ = −0.5, −0.2 and 0.1). While it may appear that the example trajectories cross against the direction of the vector field (white arrows), it is important to remember that the vector field changes over the course of the trajectory; the shown vector field is computed at the final time point in the trajectory, after the system reaches its steady state (the complete dynamics over the course of the trajectory can be seen in Movie 2). From the shown vector field, we see evidence for three fixed points of the system (Fig. 5a top row, pink circle and triangles). As the context cue increases from -0.5 to 0.0, the stable node (Fig. 5a top row pink circle) and saddle (Fig. 5a top row left pink triangle) come together and eventually coalesce at the onset of oscillations at *I*_*cc*_ = 0.18 (not shown). Recall, we had predicted this behavior from our bifurcation analysis of the reduced model: a long-period oscillation appears as a homoclinic orbit as the saddle-node bifurcation point is approached from negative to low values of context cue (Fig. 3g and Fig. 4b, red and black branches come together at the start of the green branch).

**Fig 5.**
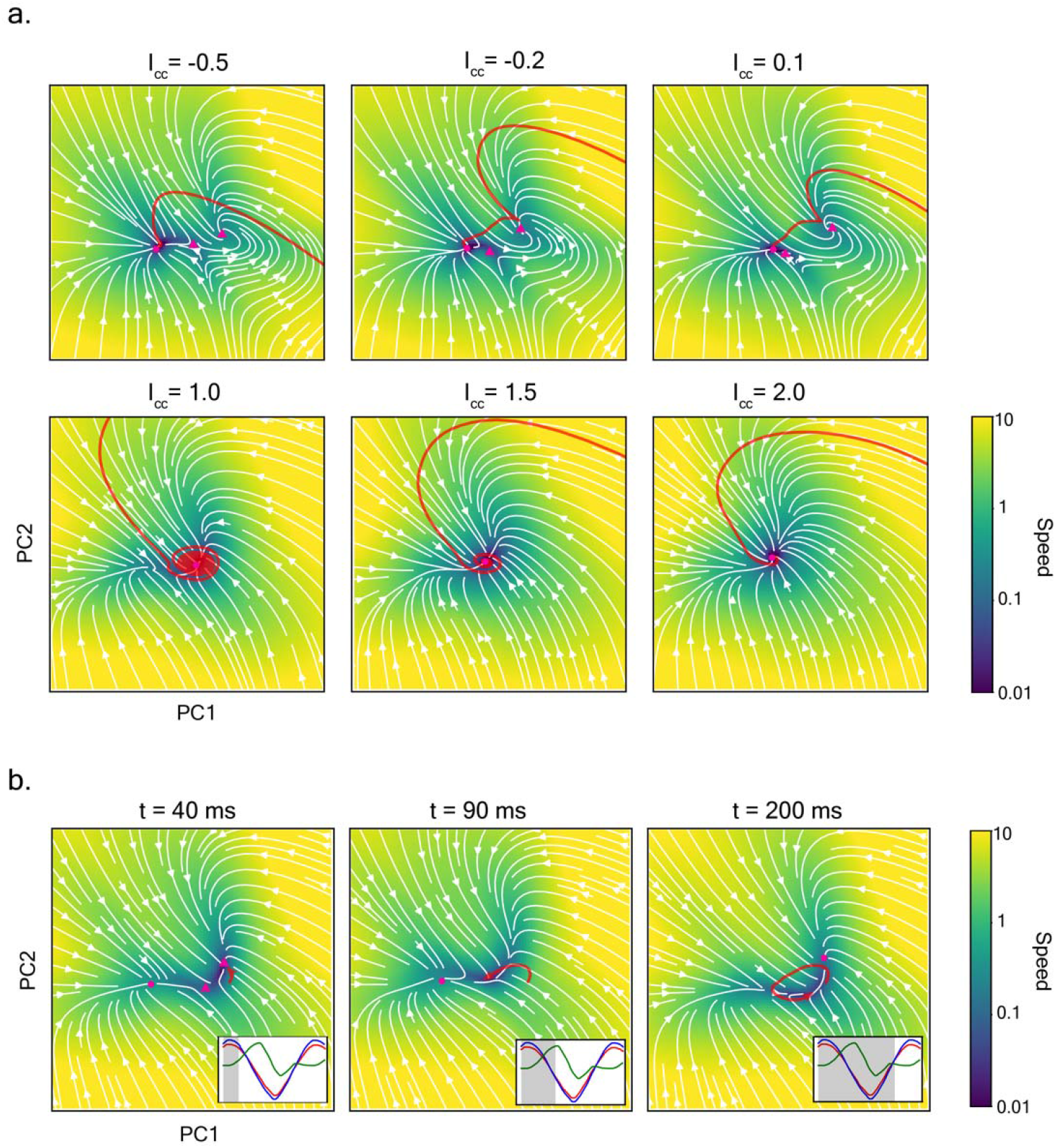
RNN dynamics agree with reduced model predictions. **a**. Two-dimensional projection of RNN dynamics for different values of context cue. In the top row, context cues *I*_*cc*_ = −0.5, −0.2 and 0.1, there is a global stable steady state attractor on the left-hand side (dark purple region) and all trajectories end up there. In the bottom row, context cues *I*_*cc*_ = 1.0, 1.5 and 2.0, the dynamics show a stable attractor again but now the trajectories spiral towards this fixed point. White arrows indicate the vector field in the projection plane; projected directions computed after the dynamics have evolved to be near steady state. Red curves indicate example trajectories. Color indicates speed with dark purple corresponding to slow speeds and bright yellow corresponding to fast speeds. Pink markers indicate fixed points deduced from flow maps with circles indicating stable fixed points and triangles indicating unstable fixed points. **b**. Two-dimensional projection of RNN dynamics for *I*_*cc*_ = 0.5 at three different time points during one cycle (period = 222 *ms*) of the oscillating trajectory. Colors are the same as in a. Pink circles and triangle indicate fixed points that are transient. Since the trajectory is oscillating, the vector field changes over time and the transient fixed points appear and disappear over the course of the oscillation. Inset shows averaged time courses for Tap-E (red), Tap-I (blue) and INT-I (green) sub-populations from the RNN with grey shading to indicate depicted time point relative to the time course.

The second row of Fig. 5a shows example trajectories for larger values of context cue, *I*_*cc*_ = 1.0, 1.5 and 2.0, projected into the same plane. The vector field for these projections is also computed once the system has reached steady state. An example trajectory (red) spirals towards the fixed point and as the context cue is increased, the fixed point becomes a stronger attractor and less spiraling is observed (Fig. 5a bottom row, compare panels from left to right). This behavior is indicative of a Hopf bifurcation in the RNN (at a context cue around *I*_*cc*_ = 0.95) and corroborates its prediction from the reduced model (Fig. 3g and Fig. 4b, for 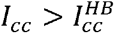.

Next, we describe the behavior of the RNN in the oscillatory regime (0.18 ≤ *I*_*cc*_ ≤ 0.95) for which the vector field changes over the course of the oscillation. We look at the vector field of an example trajectory (*I*_*cc*_ = 0.5) at three different snapshots in time (the complete dynamics can be seen in Movie 3). Figure 5b (left panel) shows the projection of an oscillating trajectory (red curve) at the time the averaged Tap-E and Tap-I time-courses are near their maximum activity (inset). Two pink circles and one pink triangle indicate transient “pseudo fixed” points of the system at the depicted point in time. These points are not fixed points of the system since they will disappear over the course of the oscillation, but they and their resulting vector fields do transiently affect the dynamics. The oscillation trajectory begins to head toward the right most circle in the snapshot. These three transient points resemble the three points of intersection of the nullclines of the reduced model (Fig. 3e magenta circles).The middle panel of Fig. 5b shows the next snapshot, when Int-I activity is at its maximum and *x, y* are decreasing (inset) and the bottom fixed point becomes a strong transient attractor and draws the trajectory toward it (for large enough *z*, the upper two fixed points in magenta disappear; see Fig. S3). This corresponds to the fixed point (magenta) seen on the bottom left of dashed nullclines in Fig. 3e. Finally, the right panel of Fig. 5b shows the example trajectory returning to its starting point and completing one oscillation cycle as the averaged Tap-E and Tap-I time courses ramp up their activity toward their maximum (inset). This snapshot of RNN behavior resembles the snapshot seen in the reduced model (Fig. 3e solid line nullclines). Notice that in Fig. 3e, the example trajectory of the reduced model, during the rising phase of *x* and *y*, heads toward that fixed point, moving just below the balance manifold (Fig. 3e, overlapping region of nullclines for *x,y* ≥0). The RNN trajectory also appears to travel along a similar manifold (purple region in Fig. 5b, right panel) toward the fixed point.

At the level of individual units, one of the signatures of a homoclinic bifurcation is an elongation of near constant activity in the firing time course as the oscillation period increases. This happens because, after the saddle and attractor coalesce, the attractor node leaves a ghost which slows trajectories that pass near to where the attractor had been. As the context cue increases, the ghost has less of an effect thereby reducing the window of slow speeds and thus decreasing the oscillation period. We see this elongation of the period of near constant activity with decreasing *I*_*cc*_ prominently in the reduced model as well as the RNN (Fig. S2). Finally, another feature of the reduced model is that the oscillation amplitude (the distance between the green curves in Fig. 3e) decreases as the Hopf bifurcation is approached – this is confirmed in the RNN (Fig. S4).

## Discussion

We developed a neurally plausible mechanism for the synchronization and continuation task (SCT) that is controllable via the context cue to oscillate over a frequency range that matches the range of beat frequencies perceived by macaques and humans. By developing a reduced model, we were able to predict and confirm the dynamical features that gave rise to oscillations in the RNN. Based on our identification of three distinct neural populations in the RNN network that are necessary for oscillations, our reduced model is comprised of an excitatory and one group of inhibitory units that form a balanced sub-network with a second group of inhibitory units that create an elongated window of inhibition that allows the system to oscillate.

The oscillation mechanism can be differentiated from previous models for rhythmic timing. Unlike entrained oscillator models^16–20^ where an endogenous brain rhythm entrains to the stimulus frequency, our model does not depend on sustained entrainment. Rather, the internally generated oscillation is tuned to the appropriate frequency through adjustment of the context cue. The mechanism that we discovered through the RNN is more closely aligned to that of tunable oscillator models such as the Beat Generator (BG) Model^21,22^ or the SAM-MPM model^23^ where the oscillation frequency adapts to match that of the stimulus through learning rules. Unlike the BG model which achieves slow oscillation by the incorporation of a slow variable, our model shows long period oscillations arising through a homoclinic bifurcation. Moreover, our model can be viewed as a novel network-based circuit model in contrast to a cellular oscillator model in the BG framework with the context cue (*I*_*cc*_) as the control parameter that sets oscillation period. Finally, other models have addressed more complex rhythms^24,25^ but have done so at the cost of losing the ability to explain the neural dynamics observed on the short timescale as we have done here. Future work could apply the current framework to more complex tasks.

With insights garnered from the RNN and reduced models, we suggest some experimental predictions that could help test our model. First, our models have two distinct sub-populations of inhibitory units: the Tap-I group that is highly synchronized to the Tap-E units and the Int-I group that shows sequential structure in the firing pattern. This architecture is consistent with previous experimental findings in other brain areas such as the visual cortex^26^ and auditory cortex^27^, where parvalbumin-expressing (PV+) inhibitory interneurons pair up with co-tuned excitatory cells as well as with somatostatin (SOM) interneurons. Oscillations are among the growing evidence for functional importance of such canonical circuitry^28,29^. A model of gamma oscillations in visual cortex is based on strong E-I balance based on PV+ inhibitory interneurons with somatostatin (SOM) interneurons playing a critical role in producing the oscillations^30^ – albeit this is for higher frequency oscillations than considered here. Taken together, these results suggest that the Tap-I units in our RNN might be mapped onto PV+ while Int-I units might be mapped onto SOM interneurons. We therefore suggest that SOM-like functional inhibition provides a mechanism for the elongated inhibition. Second, subject to neuronal subgroups being identifiable in experimental data, the firing rate time courses for this E-I pairing are expected to be highly synchronized. Third, the formulation of the reduced model allowed us to disregard the inter-tap excitatory group suggesting that these cells could be silenced without destroying the network oscillation (albeit the overall frequency may change; Fig. S3). Finally, the presence of a homoclinic bifurcation in our model suggests that as the oscillation period slows, there should be a sub-threshold plateau of near constant activity that immediately precedes firing during the “Tap” phase of activity. Another expected signature of homoclinic behavior is the lack of resonance in the autocorrelation of ITIs. In contrast, resonance would be expected for the shorter period oscillations associated with a Hopf bifurcation mechanism^31^.

In the three-variable reduced model, the variables *x* and *y* are in strong balance as exemplified by the highly synchronized time courses of *x* and *y* (Fig. 2c). This balance is reminiscent of an inhibition-stabilized-network balance operating in a regime where slow modulation of input leads to *x* and *y* both increasing or both decreasing in super-threshold activity ranges^32,33^. This behavior is unlike the noise-driven activity found near a subthreshold E-I balance state in the fluctuation-dominated regime described in spiking networks^34^.

The RNN shows sequential firing thereby achieving ITI timescales of 100 *ms* despite the utilization of 10 *ms* time constants. In an effort to retain simplicity in our three-variable model, we chose the time constant, τ_*z*_, of the Int-I group to be longer than that of Tap-E and Tap-I. We propose the interpretation that, functionally, sequential firing in the RNN (and in the neural data) serve to provide an extended duration of inhibition, acting effectively as if Int-I decays more slowly. We found little evidence of strong chain-like connectivity suggesting that our sequential structure is unlike that seen in songbirds^35^ and is more related to sequences formed from parametric gradients of excitability as seen in the hippocampus^36^ and in the posterior parietal cortex^37^. Interestingly, although more units are involved in slower frequency oscillations, we saw no increase in dimensionality (as measured by number of PCs need to explain the same level of variance) for longer sequences. In future work, it would be worthwhile to explore the effect of explicitly adding chain-like^38^ structures into the reduced model or to train a spiking network with spike-time-dependent plasticity with heterosynaptic competition on the SCT and to test if sequential activity emerges as suggested by previous computational studies ^39^.

The ease of training RNNs and their success at replicating features of neural dynamics seen in experimental data has fueled interest in uncovering the mechanisms underlying the RNN solutions to tasks^40–47^. While we do not provide a one-size-fits-all approach to reducing a high dimensional network to an interpretable circuit, we do show how close inspection of the RNN unit activity can lead to the formulation of a circuit model with distinct neuron subpopulations. In doing so, we are better able to understand the functional mechanisms that underlie activity of the hidden layers of the RNN beyond those layers simply being a black box.

By training and then dissecting an E-I RNN, we have generated a novel, neurally plausible network model of rhythmic timing. Although previous work has described the occurrence of homoclinic bifurcations in rate models for neural networks^48–51^, finding such a dynamical feature in a timing task is new. Furthermore, our model and analysis not only offers interpretations of neural dynamics previously observed in experimental data but also posits functional significance for different interneuron subtypes.

## Methods

### RNN Formulation

The RNN consists of 500 firing rate units with distinct excitatory (80%) and inhibitory (20%) populations. The output of the network was required to produce a sinusoidal response lasting for at least two seconds with peaks of the output cycles aligned with the stimulus tones. We use the peak times of the sinusoidal output as a proxy for the command signal that would be sent to execute a motor action that would be synchronized with the stimulus pulses in this task and we will refer to peak times of the output cycles as model ‘tap’ times. The network was trained on seven different frequencies: 2,3,4,5,6,7 and 8 Hz. A natural range of frequencies for musical and speech rhythms^1,52^.

The dynamics of the RNN are described by the following equations:

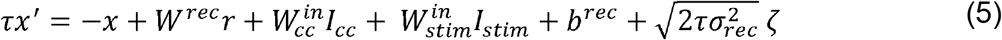

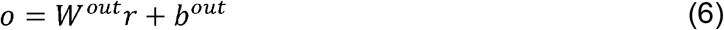

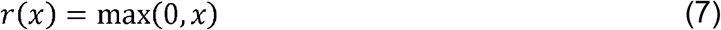

where *x* ∈ ℝ^*N*^ is activity (analogous to a mean voltage^53^) of the N (=500) RNN hidden units which are transformed into firing rates *r* via the ReLU non-linearity in equation (7). The recurrent weight matrix is given by *W*^*rec*^, the vector *b*^*rec*^ denotes the biases for each unit and *ζ*, denotes Gaussian white noise. The input weight scalars are given by 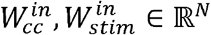. The two types of input: context cue and stimulus pulses are indicated by *I*_*cc*_ and *I*_*stim*_, respectively. The time constant of *τ*=10 *ms* was used for all units. The output (*o*) of the network is a linear combination of the firing rates with the weight matrix *W*^*rec*^ ∈ ℝ^*N*^ and a bias term *b*^*rec*^ . RNN was trained w 0.01_ ith both recurrent noise (*σ*_*rec*_ = 0.01) and input noise (*σ*_*in*_ = 0.01). Self-connections were not permitted in the network to encourage sequential firing^54^ . The quantities 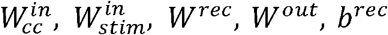, and *b*^*out*^ are ‘learned’ during training of the RNN.

### RNN Training

The RNN was trained using continuous-time dynamics – equations (5)-(7). Integration in time was by Euler’s method with a step size, dt, equal to 1 *ms*. The RNN was trained on the NYU computing cluster for 72 hours and reached an *RMSE* = 0.13.

For each of the inputs (*I*_*cc*_, *I*_*stim*_) to the network, the euler step increment at time t is modeled using:

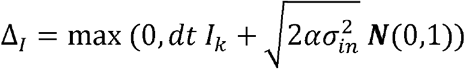

where *I*_*k*_ = *I*_*cc*_ *or I*_*k*_ = *I*_*stim*_, *α* = *dt*/*τ* and *σ*_*in*_ = 0.1 corresponds to the standard deviation of the input noise.

The network was trained using the Pycog library^55^ using stochastic gradient descent algorithm with a learning rate of 0.01. We used the mean squared error cost function between the network output (o) and target function (z):

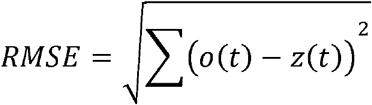

Weights ***W***^*out*^, ***W***^*in*^, ***W***^*brec*^, ***b***^*rec*^, ***b***^*out*^ were all trained. Unless otherwise specified, all other training parameter are the defaults as given in Song et al.^55^.

### Synchronization and Continuation Task

The RNN was trained with seven different pairs of inputs for *T* = 2*s* (frequencies = 2,3,4,5,6,7,8 Hz, equivalently periods = 500, 333.3, 250, 200, 166.6, 142.8, 125 ms). The inputs are of two types: *I*_*stim*_ which was provided for 1*s*, and *I*_*cc*_ which was provided for 2*s* and defined as follows:

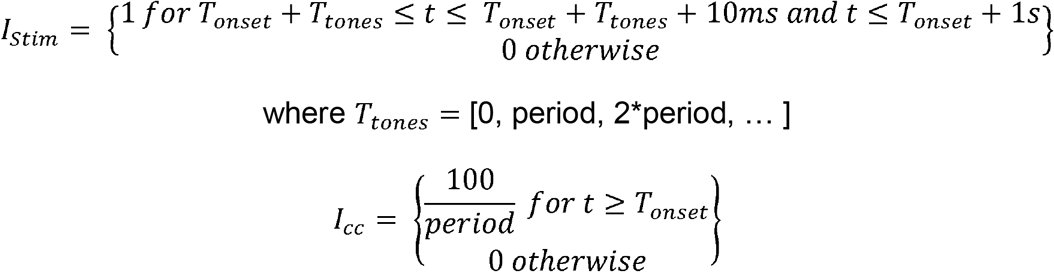

The output target was defined as

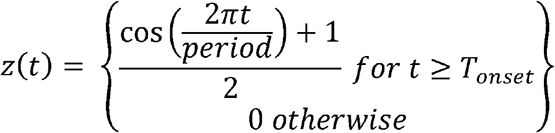

*T*_*onset*_ was random on each trial and drawn from Uniform(0,100ms).

### Generalization performance

To test how the trained RNN generalizes to previously unseen frequency inputs, we gave the trained RNN context cue inputs in the range of 0.1 to 1.1 with a step size of 0.01. RNN inter-tap intervals were computed from 50,000 *ms* of the continuation phase RNN output and each ITI was plotted in Figure 1f.

### Computation of RNN unit dynamics

Figure 1 panels **c**-**e**: Statistics of RNN trajectories are computed by first simulating the RNN with noise (*σ*_*in*_ = 0.1, *σ*_*rec*_ = 0.1) only during the continuation phase (i.e. the RNN was not provided with a stimulus pulse sequence) for 52,000 timesteps (dt = 1ms) at each *I*_*cc*_ .The first 2,000 timesteps are deleted to make sure that there are no transients and units are in their near-periodic activity patterns. Each 50,000 timestep time course is then divided into its inter-tap-intervals (ITIs) and statistics are computed over the corresponding set of ITI’s for each *I*_*cc*_.

Figure 2 panels **a, c**: RNN trajectories are simulated without noise ((*σ*_*in*_ = 0.0, *σ*_*rec*_ = 0.0) only during the continuation phase. The first 2,000 *ms* of a time course are dropped to make sure transients have passed and units are in their periodic activity states. Each unit is normalized so that the firing activity falls between 0 and 1. The normalized units are sorted by time of peak firing (when firing activity is at 1) during their first ITI after the 2,000 *ms* of dropped time course (excitatory and inhibitory are sorted separately). Units that hit their max firing rate with a phase of within 0.2 of the model “tap” times are labeled ‘tap’ units and the remaining units are labeled inter-tap units.

### Analysis of RNN dynamics

The projection plane for the analysis of RNN dynamics was kept constant for all panels in Fig. 4. This plane was selected by first simulating one hundred T=500 *ms* noise-free trajectories of RNN units with context cues from the range of -1 to 0.2. On each simulated trajectory, the initial conditions for the hidden variables were selected so that the units that were grouped into Tap excitatory and Tap inhibitory units had the same value which was randomly chosen on each trial from the Uniform (0,1) distribution. All other units were individually randomly initialized in Uniform (-1,1). These trajectories were then concatenated together into a matrix of size T * 100 x N where T is the number of timesteps computed for each trajectory and N is the number of units in the RNN. Principal component analysis was performed on this matrix after it was normalized for each unit. The plane spanned by the first two principal components was selected for the projection. For noise-free trajectories, the top 2 PCs accounted for 80 percent of the variance and the top 3 PCs accounted for 85 percent of the variance.

This choice of initial conditions for these trajectories was motivated by the observation that in the reduced model, several fixed points lie on the balance manifold (in the region where x and y nullclines are almost overlapping for *x, y* ≥0). Initializing trajectories at different points along this manifold meant that the resulting trajectories should have their flows affected by these fixed points or the resulting strong vector field near to the balance manifold if the reduced model is a good proxy for the dynamics in the full RNN. PCA would then represent the directions in state-space that capture these dynamics as well. Indeed, we find that initializing points in the RNN near the predicted balance-manifold allowed PCA to find a plane capturing the predicted unstable fixed points. Initializing trajectories randomly in the full 500-dimensional space, in contrast, will capture the stable fixed point since all trajectories head toward the attractor but is unlikely to create trajectories that pass close enough to the balance manifold since the region of space that it influences is relatively small. In agreement with this, we found that initializing trajectories randomly (instead of around the balance manifold) and then performing PCA consistently only captured the stable fixed point but not the predicted unstable fixed points. Although we don’t know with certainty where the balance manifold lies in the full RNN, initializing all units that were grouped into either tap excitatory or tap inhibitory units to the same (random) value for each simulated trajectory worked well. It was important that we capture the plane with the fixed points on the balance manifold because, according to the reduced model, these fixed points are involved in the homoclinic bifurcation and we wanted to investigate if this was the case in the RNN as well.

Having defined the projection plane, the vector field was computed by first defining a vector, 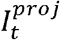, for each point in the projection plane to take the values in the plane for the first two components and filling the remaining components with their corresponding value at time t for a given example trajectory. This vector was then projected back to the full RNN state space, 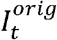, and simulated one step forward using the RNN to get, 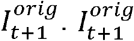. was then projected back into the space defined by the principal components to get 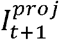 and the vector flow at point 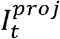 was given by the direction of 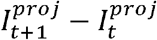. Speed is the *L*^2^ norm of the RNN hidden unit activity, *dx*_*i*_ */ dt*, computed similarly with dt=1ms:

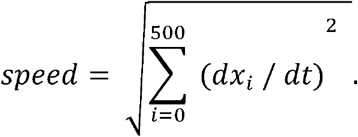

### Reduced model

The three (equations (1)-(4)) and two variable reduced models were discretized for simulations using Euler method with a timestep of *dt* = 1*ms*. Reducing the timestep to *dt* = 0.1 *ms* did not change the results.

Weights for the reduced model were taken to be the between sub-population averages from the RNN except for *τ*_*z*_, *W*_*zy*_, *W*_*zz*_ . As discussed in the text, the *τ*_*z*_ was increased to have oscillation periods in the reduced models more closely match up with the RNN. The exact value of *τ*_*z*_ does not affect the key findings as long as it is set to be in the oscillatory regime (Fig. 3h). The RNN averaged connectivity weights for *W*_*zy*_ and *W*_*zz*_ were such that the reduced model did not show oscillations and were adjusted by hand to put the model into the oscillatory regime.

### Bifurcation Analysis

All bifurcation analysis was done using XPPAUT^56^ and the resulting data was imported into python to create the figures.

## Acknowledgements

K.Z: *This work is supported by the Google Ph*.*D. Research Fellowship*.

A.B.: *This material is based upon work supported by the National Science Foundation under Grant No. DMS-1929284 while the author was in residence at the Institute for Computational and Experimental Research in Mathematics in Providence, RI, during the* Math + Neuroscience: Strengthening the Interplay Between Theory and Mathematics *program*.

## Supplementary Figures

**Fig S1.**
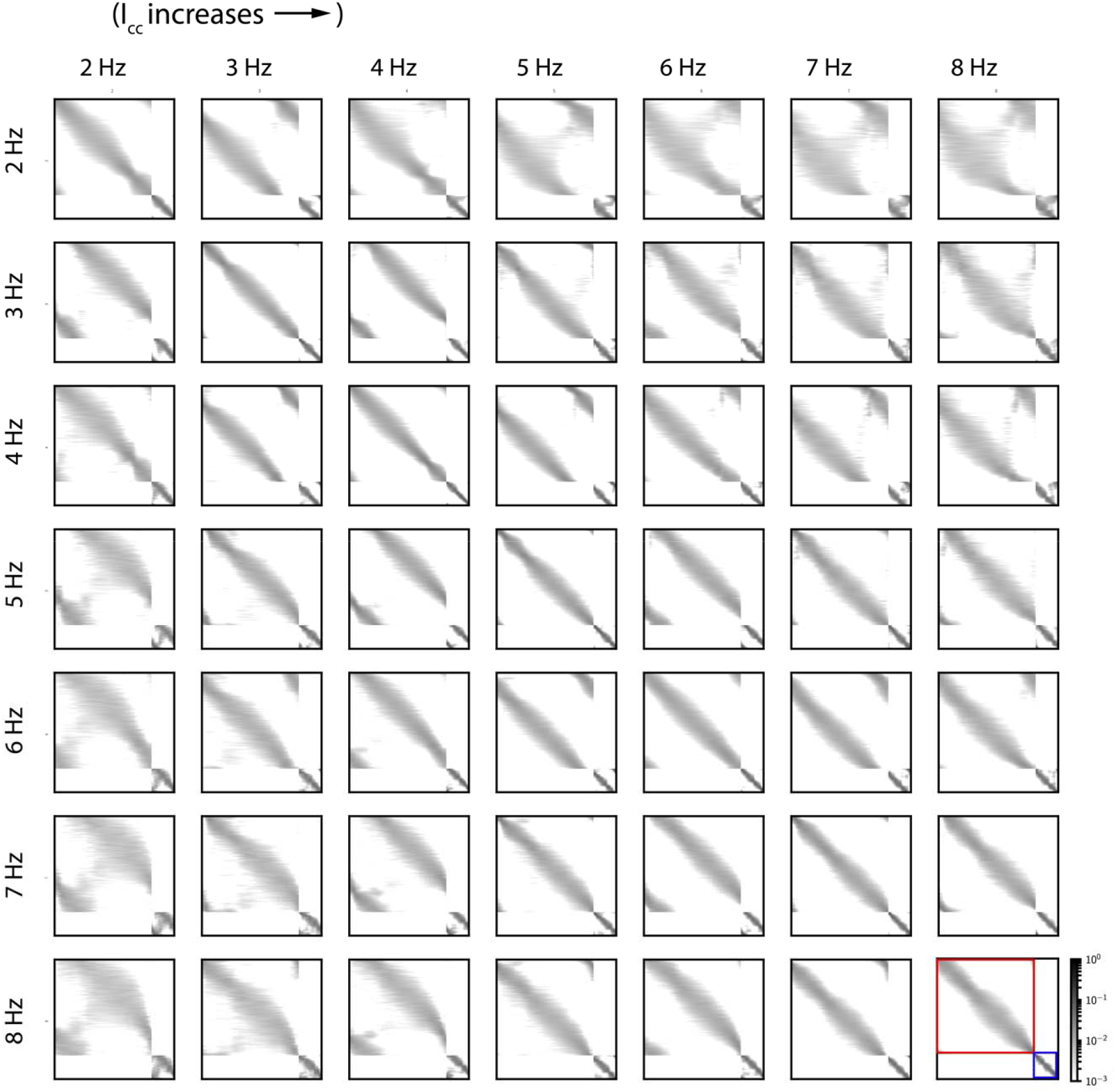
Ordering of unit sequences is highly preserved for nearby frequencies. Comparing the sequence of units after they have been sorted according to their time of peak firing rate for one stimulus frequency versus another reveals that while neurons do change their position in the firing sequence, they generally move to a firing position that is relatively close to their previous position. For each trial of a given stimulus frequency, neurons are sorted according to the time of their peak firing during one ITI of the continuation phase. Grey intensity at position (i,j) of panel (n,m) indicates the probability that a unit that fired at position i according to the sorting of frequency n will fire at position j according to sorting of frequency m. Excitatory and inhibitory units are sorted separately: for each panel, the upper block corresponds to the excitatory units while the lower block corresponds to the inhibitory units (as illustrated by the red and blue squares of panel (7,7), respectively). The panels on the diagonal indicate how much variability there is in sequences for the same stimulus and thus indicates how much the firing order changes due to noise. The ordering of the inhibitory units seems much more preserved as indicated by the narrower grey cloud along the diagonal for the inhibitory block vs. the excitatory block. There is a small amount of shifting of units between the end and the beginning of each sequence (indicated by the cloud of grey in off-diagonal corners of the excitatory block) but this is to be expected since noise can drive units to fire earlier or later than the ‘tap’ thus affecting their position in the ITI. Comparing the 2 Hz row and 3 Hz column (this corresponds to increasing *I*_*cc*_), the grey cloud along the diagonal has shifted downward which suggests that units that fired later in the sequence under 2 Hz now appear earlier in the sequence for 3 Hz. Meanwhile, the probability that a unit that appeared in the middle of the sequence for the 2 Hz sequence would now appear at the beginning or end of the sequence for the 3 Hz sequence is close to zero – this is indicated by the presence of white off of the diagonal in each panel. The opposite pattern occurs when *I*_*cc*_ decreases; units that fired earlier in the sequence for 3 Hz now fire later in the sequence for 2 Hz (3 Hz row, 2 Hz column). Probabilities are computed over 10,000 trials of each frequency.

**Fig S2.**
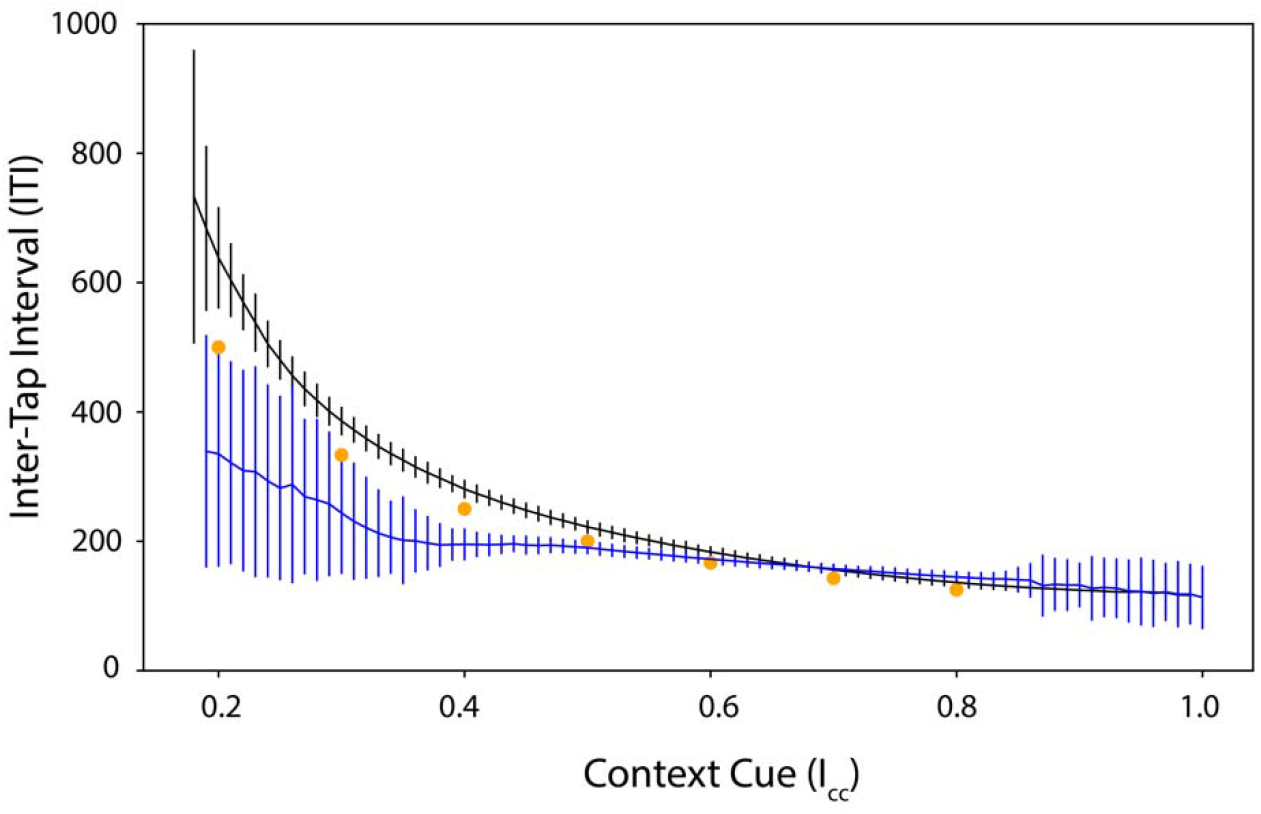
Oscillations persist in RNN after silencing inter-tap excitatory group. Simulation of RNN ITIs for varying levels of context cue before (grey) and after (blue) silencing all of the Int-E units. Vertical lines indicate mean ± std. dev. of the ITIs of the model output during 50,000 *ms* of simulated continuation phase for each context input. Yellow circles indicate trained points. Silencing the Tap-E population has the effect of speeding up and increasing the variability of the oscillations for low frequencies.

**Fig S3:**
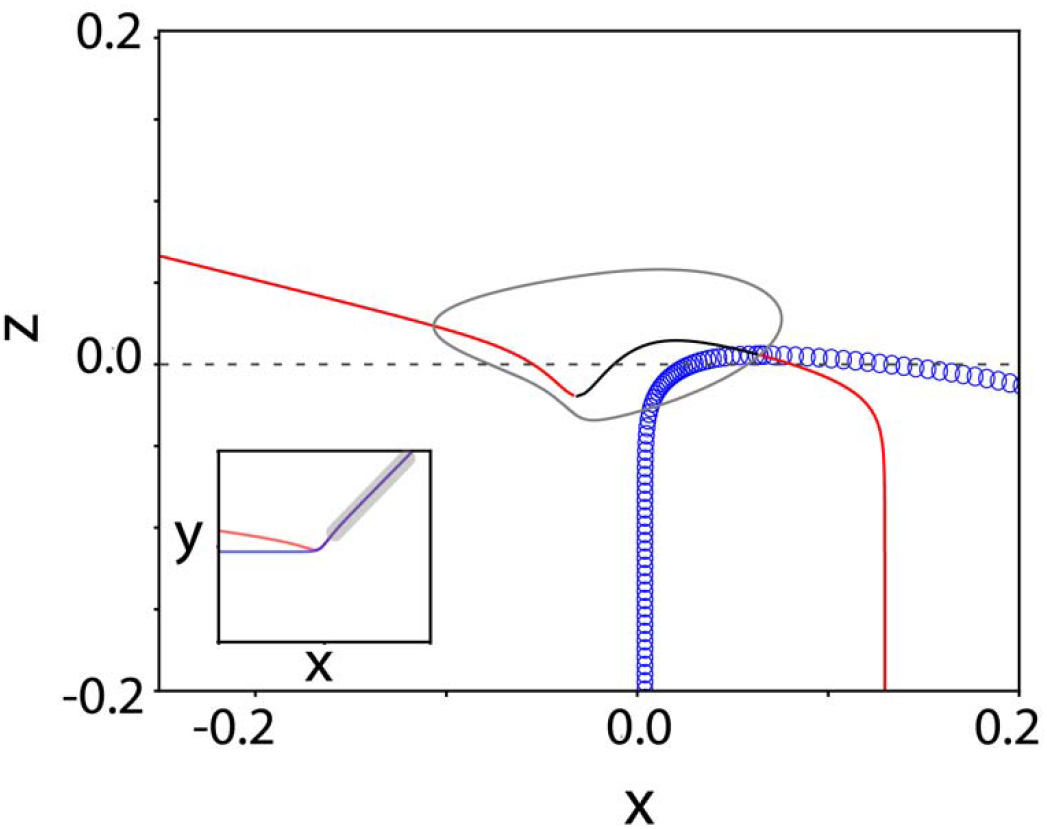
Bifurcation diagram with z as a control parameter of x-y subsystem of 3-variable reduced model at *I*_*cc*_ = 0.5. For fixed levels of *z*, the *x* −*y* **sub**system shows stable steady states (red lines), unstable steady states (black line) and unstable oscillations (blue open circles). The left-hand stable branch corresponds to a strong attractor. The right-hand stable branch is also an attractor but is surrounded by an unstable orbit so trajectories cannot get to the attracting state unless they start near it. Limit cycle (stable) trajectory for the 3-variable model (black closed curve) is superimposed, showing that the left-branch is tracked closely during the INT-phase. The inset shows a snapshot of the *x* −*y* phase plane with grey indicating the very small region that is surrounded by the unstable orbit when *z =* 0 (dashed line).

**Fig S4.**
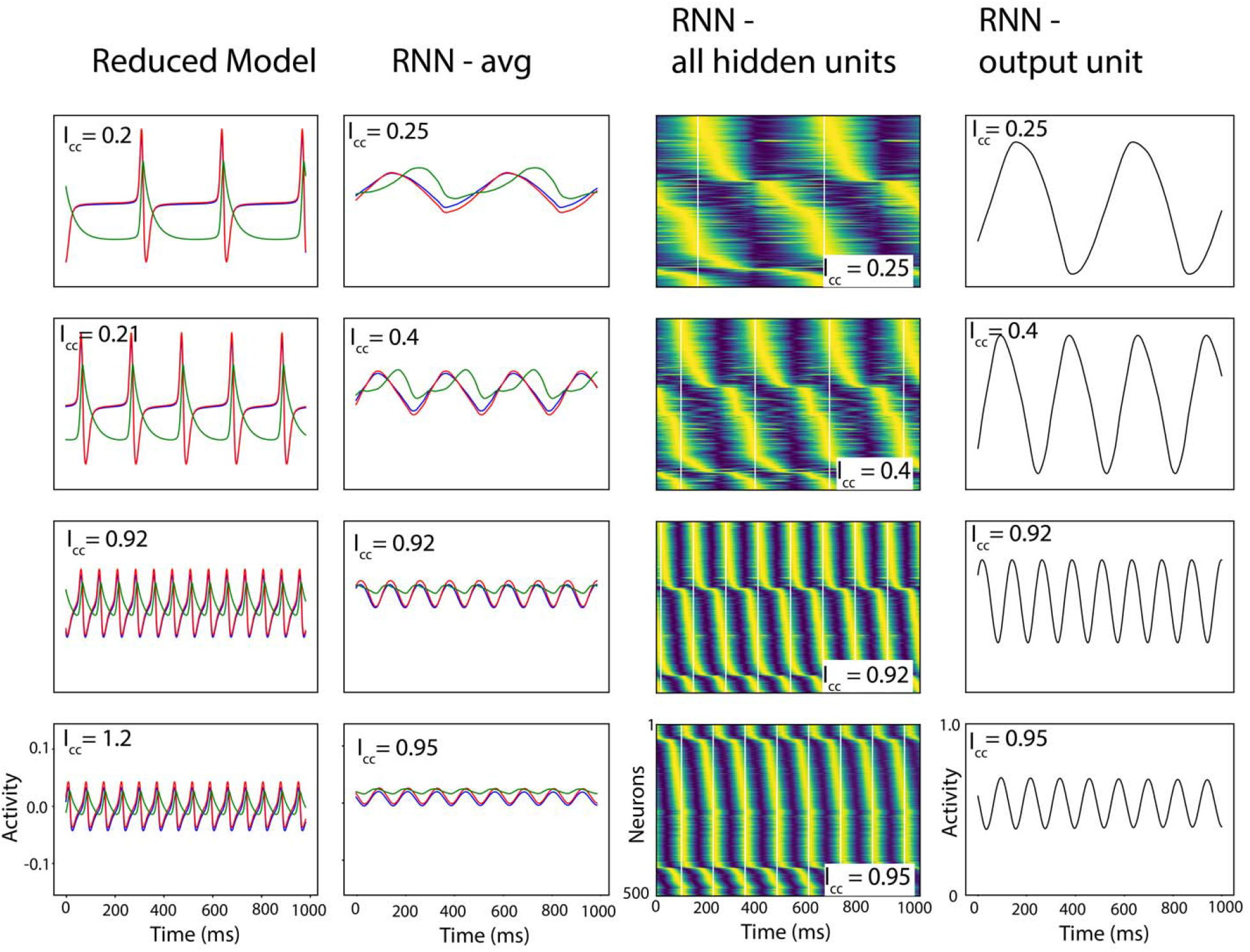
Oscillation time courses for RNN and reduced model. First column shows the reduced model time courses for different levels of context cue (indicated in panel). Red, blue and green traces show the activity for Tap-E (*x*), Tap-I(*y*) and Int-I (*z*) variables, respectively. Note that the as the context cue decreases (moving up the column), the period of activity where the and variables are near zero increases – this is an indication of the homoclinic bifurcation. The second column shows the firing activity time courses for RNN units averaged by group. Red, blue, and green correspond to the average firing activity of hidden units in the Tap-E, Tap-I and Int-I subpopulations. Note that the amplitude of the oscillations decreases as context cue increases (moving down the column) – this is indicative of approaching the Hopf Bifurcation at a high level of context cue. Third column shows the normalized activities for all hidden units in the RNN with yellow colors indicating the maximum activity for that unit and dark blue indicating the lowest activity for that unit. Excitatory and inhibitory units are sorted separately: excitatory units are shown in the first 400 rows and inhibitory units are shown in the bottom 100 rows for each panel. White vertical lines indicate model “tap” times. Note that individual units (rows in subpanels) show an elongated period of low firing (dark blue) as context cue is decreased (moving up the figure column) – this is suggestive of the homoclinic bifurcation in the RNN. Fourth column shows the activity of the RNN output unit. All depicted trajectories are simulated without noise. The reduced model trajectories (first column) have notable different qualities from the RNN sub-population averaged timecourses (second column): the reduced model trajectories are substantially more negative, show a more pronounced period of near constant activity and have a more pronounced spike-event.

**Movie1: Link:**https://drive.google.com/file/d/1OixLeOv_5paX1SQV1gSy6ejStnE0ie3Y/view?usp=drive_link

Phase plane projections for *x* −*y* (left panel), *x* −*z*(middle panel) and time course of activity for 3-variable reduced model. Nullclines are shown in red, blue and green for *x,y* and *z* variables, respectively. Black trace in right and middle panels shows example trajectory projected onto the corresponding plane. Right panel shows time course of *x* (red), *y* (blue) and *z* (green) variables during one oscillation cycle. In the left panel, the oscillation hugs the E-I balance manifold (overlapping *x, y* nullclines in *x, y* ≥ region) until *z* becomes sufficiently large so that the fixed point on the bottom left of the appears and acts like a strong attractor. As is reduced, this point disappears and the trajectory heads back to the *x, y* ≥ region.

**Movie 2:**

**Link:https://drive.google.com/file/d/1bLTUIvc8iaE7-dzc4ErxCo8NzUyyzWct/view?usp=drive_link**

Projection of RNN activity for *I*_*cc*_= −0.2, Where the RNN converges to a stable fixed point, into the plane spanned by the first two principal components. Example trajectory is shown in black. Vector fields at each time point are indicated in white and speed is indicated by the color. Color map is the same as in Fig. 5. Once the system reaches the steady state, three fixed points can be seen (dark blue regions) but on the left-most one is stable (which is the one that the trajectory ends up at).

**Movie 3**: **Link:https://drive.google.com/file/d/1Trfs3mHZuEieAwkSRBkQ_n57_dvcRGsZ/view?usp=drive_link**

Projection of RNN activity for an example time course at *I*_*cc*_ =0.5, when the RNN shows oscillations, into the plane spanned by the first two principal components. Example trajectory in red. Tone of red indicates activity of average inter-tap population activity with white indicating low firing and dark red indicating high firing. Vector fields at each time point are indicated in white and speed is indicated by the color. Color map is the same as in Fig. 5. The oscillation begins heading in the direction of the upper, right fixed point then turns around to head toward the bottom-left fixed point which acts as a strong attractor (when the *z* activity is very high as indicated by the dark red shading of the trajectory). Finally, the trajectory turns around and heads back toward its starting point along the slow manifold (dark blue).

## Notes

### Competing Interest Statement

The authors have declared no competing interest.

## References

1. Repp, B. H. Sensorimotor synchronization: A review of the tapping literature. Psychonomic Bulletin and Review 12, 969–992 (2005).

2. Repp, B. H. & Su, Y. H. Sensorimotor synchronization: A review of recent research (2006-2012). Psychon. Bull. Rev. 20, 403–452 (2013).

3. Gámez, J., Mendoza, G., Prado, L., Betancourt, A. & Merchant, H. The amplitude in periodic neural state trajectories underlies the tempo of rhythmic tapping. PLOS Biol. 17, e3000054 (2019).

4. Jazayeri, M., Merchant, H., Gracía-Garibay, O. & Malagón, A. M. Keeping time and rhythm by replaying a sensory-motor engram 1 2 3 Victor de Lafuente. doi:10.1101/2022.01.03.474812

5. Patel, A. D. Vocal learning as a preadaptation for the evolution of human beat perception and synchronization. Philos. Trans. R. Soc. B Biol. Sci. 376, (2021).

6. Betancourt, A., Pérez, O., Gámez, J., Mendoza, G. & Merchant, H. Amodal population clock in the primate medial premotor system for rhythmic tapping. Cell Rep. 42, 113234 (2023).

7. Large, E. W. et al. Dynamic models for musical rhythm perception and coordination. Front. Comput. Neurosci. 17, (2023).

8. Russo, A. A. et al. Motor Cortex Embeds Muscle-like Commands in an Article Motor Cortex Embeds Muscle-like Commands in an Untangled Population Response. Neuron 97, 953-966.e8 (2018).

9. Mante, V., Sussillo, D., Shenoy, K. V. & Newsome, W. T. Context-dependent computation by recurrent dynamics in prefrontal cortex. Nat. 2013 5037474 503, 78–84 (2013).

10. Goudar, V. & Buonomano, D. V. Encoding sensory and motor patterns as time-invariant trajectories in recurrent neural networks. Elife 7, (2018).

11. Sohn, H., Narain, D., Meirhaeghe, N. & Jazayeri, M. Bayesian Computation through Cortical Latent Dynamics. Neuron 103, 934–947.e5 (2019).

12. Sussillo, D., Churchland, M. M., Kaufman, M. T. & Shenoy, K. V. A neural network that finds a naturalistic solution for the production of muscle activity. Nat. Neurosci. 18, (2015).

13. Beiran, M., Meirhaeghe, N., Sohn, H., Jazayeri, M. & Ostojic, S. Parametric control of flexible timing through low-dimensional neural manifolds. SSRN Electron. J. 111, 739–753.e8 (2021).

14. Zhou, S., Masmanidis, S. C. & Buonomano, D. V. Encoding time in neural dynamic regimes with distinct computational tradeoffs. PLOS Comput. Biol. 18, e1009271 (2022).

15. Hardy, N. F. & Buonomano, D. V. Encoding time in feed forward trajectories of a recurrent neural network model. Neural Computation 30, 378–396 (2018).

16. Kim, J. C. & Large, E. W. Signal Processing in Periodically Forced Gradient Frequency Neural Networks. Front. Comput. Neurosci. 9, 152 (2015).

17. Kim, J. C. & Large, E. W. Multifrequency Hebbian plasticity in coupled neural oscillators. Biol. Cybern. 115, 43–57 (2021).

18. Tichko, P., Kim, J. C. & Large, E. W. Bouncing the network: A dynamical systems model of auditory–vestibular interactions underlying infants’ perception of musical rhythm. Dev. Sci. 24, e13103 (2021).

19. Large, E. W., Herrera, J. A. & Velasco, M. J. Neural networks for beat perception in musical rhythm. Front. Syst. Neurosci. 9, 159 (2015).

20. Large, E. W., Almonte, F. V & Velasco, M. J. A canonical model for gradient frequency neural networks. Phys. D Nonlinear Phenom. 239, 905–911 (2010).

21. Byrne, Á., Rinzel, J. & Bose, A. Order-indeterminant event-based maps for learning a beat. 1–14 (2020). doi:10.48550/arXiv.2005.06425

22. Bose, A., Byrne, Á. & Rinzel, J. A neuromechanistic model for rhythmic beat generation. PLOS Comput. Biol. 15, e1006450 (2019).

23. Egger, S. W., Le, N. M. & Jazayeri, M. A neural circuit model for human sensorimotor timing. Nat. Commun. 11, 3933 (2020).

24. Calderon, C. B., Verguts, T. & Frank, M. J. Thunderstruck: The ACDC model of flexible sequences and rhythms in recurrent neural circuits. PLOS Comput. Biol. 18, e1009854 (2022).

25. Cannon, J. PIPPET: A Bayesian framework for generalized entrainment to stochastic rhythms. bioRxiv 2020.11.05.369603 (2020). doi:10.1101/2020.11.05.369603

26. Znamenskiy, P. et al. Functional specificity of recurrent inhibition in visual cortex. Neuron 112, 991-1000.e8 (2024).

27. Li, L. Y. et al. A Feedforward Inhibitory Circuit Mediates Lateral Refinement of Sensory Representation in Upper Layer 2/3 of Mouse Primary Auditory Cortex. J. Neurosci. 34, 13670–13683 (2014).

28. Bos, H., Oswald, A. M. & Doiron, B. Untangling stability and gain modulation in cortical circuits with multiple interneuron classes. bioRxiv 1–30 (2024). doi:10.1101/2020.06.15.148114

29. Kumar, M. et al. Cell-type-specific plasticity of inhibitory interneurons in the rehabilitation of auditory cortex after peripheral damage. Nat. Commun. 2023 141 14, 1–23 (2023).

30. Veit, J., Hakim, R., Jadi, M. P., Sejnowski, T. J. & Adesnik, H. Cortical gamma band synchronization through somatostatin interneurons. Nat. Neurosci. 20, 951–959 (2017).

31. Tateno, T., Harsch, A. & Robinson, H. P. C. Threshold firing frequency-current relationships of neurons in rat somatosensory cortex: Type 1 and type 2 dynamics. J. Neurophysiol. 92, 2283–2294 (2004).

32. Tsodyks, M. V., Skaggs, W. E., Sejnowski, T. J. & McNaughton, B. L. Paradoxical Effects of External Modulation of Inhibitory Interneurons. J. Neurosci. 17, 4382–4388 (1997).

33. Murphy, B. K. & Miller, K. D. Balanced Amplification: A New Mechanism of Selective Amplification of Neural Activity Patterns. Neuron 61, 635–648 (2009).

34. Van Vreeswijk, C. & Sompolinsky, H. Chaos in neuronal networks with balanced excitatory and inhibitory activity. Science (80-.). 274, 1724–1726 (1996).

35. Long, M. A., Jin, D. Z. & Fee, M. S. Support for a synaptic chain model of neuronal sequence generation. Nature 468, 394–399 (2010).

36. Itskov, V., Curto, C., Pastalkova, E. & Buzsáki, G. Cell assembly sequences arising from spike threshold adaptation keep track of time in the hippocampus. J. Neurosci. 31, 2828–2834 (2011).

37. Rajan, K., Harvey, C. D. D. & Tank, D. W. W. Recurrent Network Models of Sequence Generation and Memory. Neuron 90, 128–142 (2016).

38. Zemlianova, K., Bose, A. & Rinzel, J. A biophysical counting mechanism for keeping time. Biol. Cybern. 116, 205–218 (2022).

39. Fiete, I. R., Senn, W., Wang, C. Z. H. & Hahnloser, R. H. R. Spike-Time-Dependent Plasticity and Heterosynaptic Competition Organize Networks to Produce Long Scale-Free Sequences of Neural Activity. Neuron 65, 563–576 (2010).

40. Langdon, C. & Engel, T. A. Latent circuit inference from heterogeneous neural responses during cognitive tasks. bioRxiv 2022.01.23.477431 (2022).

41. Mastrogiuseppe, F. & Ostojic, S. Linking Connectivity, Dynamics, and Computations in Low-Rank Recurrent Neural Networks. Neuron 99, 609–623.e29 (2018).

42. Dubreuil, A., Valente, A., Beiran, M., Mastrogiuseppe, F. & Ostojic, S. The role of population structure in computations through neural dynamics. Nat. Neurosci. 25, 783–794 (2022).

43. Langdon, C., Genkin, M. & Engel, T. A. A unifying perspective on neural manifolds and circuits for cognition. Nat. Rev. Neurosci. 24, 363–377 (2023).

44. Schaeffer, R., Khona, M., Meshulam, L. & Fiete, I. R. Reverse-engineering recurrent neural network solutions to a hierarchical inference task for mice. in Advances in Neural Information Processing Systems 2020-Decem, (2020).

45. Barak, O., Sussillo, D., Romo, R., Tsodyks, M. & Abbott, L. F. From fixed points to chaos: Three models of delayed discrimination. Prog. Neurobiol. 103, 214–222 (2013).

46. Maheswaranathan, N., Williams, A. H., Golub, M. D., Ganguli, S. & Sussillo, D. Reverse engineering recurrent networks for sentiment classification reveals line attractor dynamics. Adv. Neural Inf. Process. Syst. 32, 15696 (2019).

47. Sussillo, D. & Barak, O. Opening the Black Box: Low-Dimensional Dynamics in High-Dimensional Recurrent Neural Networks. Neural Comput. 25, 626–649 (2013).

48. Cowan, J. D., Neuman, J. & van Drongelen, W. Wilson–Cowan Equations for Neocortical Dynamics. J. Math. Neurosci. 6, 1–24 (2016).

49. Borisyuk, R. M. & Kirillov, A. B. Bifurcation analysis of a neural network model. Biol. Cybern. 66, 319–325 (1992).

50. Ermentrout, G. B. & Cowan, J. D. Temporal oscillations in neuronal nets. J. Math. Biol. 7, 265–280 (1979).

51. Hoppensteadt, F. C. & Izhikevich, E. M. Weakly Connected Neural Networks. 126, (1997).

52. Patel, A. D. & Iversen, J. R. The evolutionary neuroscience of musical beat perception: the Action Simulation for Auditory Prediction (ASAP) hypothesis. Front. Syst. Neurosci. 8, (2014).

53. Ermentrout, G. B. & Terman, D. H. Mathematical Foundations of Neuroscience. Interdisciplinary Applied Mathematics 35, (Springer New York, 2010).

54. Orhan, A. E. & Ma, W. J. A diverse range of factors affect the nature of neural representations underlying short-term memory. Nat. Neurosci. 22, 275–283 (2019).

55. Song, H. F., Yang, G. R. & Wang, X. J. Training Excitatory-Inhibitory Recurrent Neural Networks for Cognitive Tasks: A Simple and Flexible Framework. PLOS Comput. Biol. 12, e1004792 (2016).

56. Ermentrout, B. Simulating, Analyzing, and Animating Dynamical Systems. A guide to XPPAUT for researchers and students, SIAM (Society for Industrial and Applied Mathematics, 2002). doi:10.1137/1.9780898718195

